# Nuclear and Plastid DNA Sequence-based Molecular Phylogeography of *Salvadora oleoides* (Salvadoraceae) in Punjab, India

**DOI:** 10.1101/050518

**Authors:** Felix Bast, Navreet Kaur

## Abstract

*Salvadora oleiodes* is a tropical tree species belonging to the little-known family Salvadoraceae and distributed in the arid regions of Africa and Asia. Aims of our study were to trace the microevolutionary legacy of this tree species with the help of sequence-based multi-local phylogeography and to find the comparative placement of family Salvadoraceae within angiosperm clade malvids. A total 20 geographical isolates were collected from different regions of North India, covering a major part of its species range within the Indian Subcontinent. Sequence data from nuclear-encoded Internal Transcribed Spacer region (ITS1-5.8S-ITS2) and plastid-encoded *trnL-F* spacer region, were generated for this species for the first time in the world. ITS-based Bayesian phylogeographic analysis revealed the existence of four clades while *trnL-F* spacer based Bayesian analysis revealed one clade for this species distributed in the Indian subcontinent. Between these two loci, ITS revealed more distinct phylogeographic clades, indicating the phylogeographic utility of this locus for the systematics of Salvadoraceae. Phylogenetic analyzes based on *trnL-F* spacer suggested a synonymy of this species with *Salvadora angustifolia*. Maximum Likelihood gene tree based on ITS sequence data revealed that Salvadoraceae belongs to Sapindales rather than Brassicales. However, in the gene tree based on *trnL-F* spacer sequence, this family clustered within Brassicales. An evolutionary congruence of *S. oleoides* isolates across its range in North India is revealed in this study. Given the conflicting results on the relative placement of Salvadoraceae in Brassicales and Sapindales, the need for further phylogenetic analyses of malvids using supermatrix approach is highlighted.

## Introduction

*Salvadora oleoides* Decne is a tropical tree-shrub species belong to the little-known family Salvadoraceae Lindl. This tree species has opposite individual undivided leaves, imbricated corolla lobes, tertramerous flowers with four stamens, and basally-fused connate petals. Although the natural range of this species extends from sub-Saharan Africa up to the island of Java, Indonesia, this species is under the stress of habitat destruction due to widespread deforestation and over-exploitation. Natural habitats of this tree, tropical thorn forests, are endangered by a variety of human activities, especially expansion of agricultural fields. While neither the official IUCN Red List, nor the recent estimate of threatened tree species worldwide (Oldfield et al. 1998) ascertain the conservation status of *S. oleoides*, there are some indications that this species is threatened (Khan et al. 2011; Khan 1996). In India, the tree is found in the isolated thorn forests of the arid regions of North India, especially in and around the Thar Desert.

As per the latest APG III system, the family Salvadoraceae is recognized in cabbage order Brassicales (Bremer et al. 2009). This family consists of poorly understood genera *Salvadora*, *Azima*, and *Dobera*. Similar to the other members of Brassicales, members of Salvadoraceae produces a pungent phytochemical, mustard oil glucosinolate. Production of this chemical is thought to be a synapomorphic character of the entire crown group of Brassicales (Rodman et al. 1998). Since time immemorial plants of genus *Salvadora* had been used in traditional Ayurvedic medicine for a wide variety of purported health benefits; including hypoglycemic and hypolipidemic (Yadav et al. 2008), antimicrobial (Kumar et al. 2012) and as an antibacterial toothbrush. A recent analysis of the feces of a critically endangered Lemur species endemic to Madagascar, golden-crowned Sifaka (*Propithecus tattersale*), revealed that a major portion of its diet was encompassed of the leaves of *Salvadora angustifolia* (Quéméré et al. 2013), exposing a small facet of its hidden ecological importance.

Despite its ecological, conservational and commercial importance, systematics and taxonomy of this tropical tree species have neither been analyzed using molecular systematics, nor its evolutionary legacy been traced using sequence-based phylogenetic approach. For example, the phylogenetic position of *Salvadora oleoides* within higher taxonomic hierarchies, such as Brassicales, malvids or rosids, had never been assessed to date. Prior to this study, no nucleotide sequence data had been available for this species in the GenBank database. Till date, there are only two reports available that analyzed relative position of genus *Salvadora* within malvids. The first study was based on chloroplast encoded *rbcL* gene and nuclear-encoded 18S sequences, and the study revealed phylogenetic affinity of this genus with Brassicales/Capparales (Rodman et al. 1998). The second study was based on *rbcL* gene and that too corroborated phylogenetic affinity of genus *Salvadora* within Brasicales (Savolainen et al. 2000). Genetic heterogeneity of *Salvadora oleoides* from India had previously been assessed by RAPD method and concluded a very high genetic heterogeneity, with percent polymorphism at 90.09% (Yadav 2014). Isozyme electrophoresis-based analysis also revealed a high genetic variation (H_T_ = 0.249) for this species distributed in North India (Saini and Yadav 2013).

Primary objective of this study was to assess the number, phylogenetic relationships and geographic distributions of evolutionary lineages of *Salvadora oleoides* in North India‐ a first such phylogeographic assessment of a member of family Salvadoraceae throughout the world. We were especially motivated by the pithy dictum famously declared by Croizat (Croizat 1964), that the earth and life are co-evolving. Given that the natural range of this species in Indian Subcontinent lies along the North-Western fringe of Indian tectonic plate, we were interested whether biogeographic processes akin to the Gondwanaland and Indian plate contributed in the distribution of this species. For example, species-range expansion through geodispersion phenomena during immediate aftermath of the collision of Indian plate to the Eurasian plate in Eocene epoch of Cenozoic around 50 million years ago, or vicariance events happened subsequently after the formation of the Himalayas, might have shaped the current patterns of the distribution of genetic diversity and population differentiation within species. Brassicales is believed to have originated either in Turkey or Irano-Turanian region and diversified 85 million years ago, prior to the collision of the Indian plate to the Eurasian Plate (Franzke et al. 2011). Given that conservation status of *Salvadora oleoides* is much contested amidst recommendations to classify it as a threatened species, a high-resolution assessment of genetic variation using DNA barcoding will be of paramount importance in its conservation genetics scenario as well. This is because extent and distribution of intraspecific variation is a reliable proxy for determining evolutionary potential and long-term survival of tree species (Holsinger and Gottlieb 1991). Our second objective had been to resolve the phylogenetic placement of this species and family Salvadoraceae within malvids. In many schemes of phylogenetic systematics Salvadoraceae had been considered as part of orders Brassicales/Capparales (Bremer et al. 2009; Rodman et al. 1998), Sapindales (Engler and Diels 1936), Salvadorales (Dahlgren 1989), Oleales (Goldberg 1986), Celastrales (Takhtadzhian 1997) and even as an “outsider”, *incerta sedis* (Thorne 1992). As nucleo-ribosomal Internal Transcribed Spacers 1 and 2 are the most widely covered DNA barcode for angiosperms at the NCBI GenBank (Kress et al. 2005) and as sequence information at this locus is not yet available for Salvadoraceae, we were interested to generate ITS sequence data and test various phylogenetic hypotheses on the placement of this family within malvids. In addition to ITS, we also employed plastid-encoded *trnL-F* spacer region, as no DNA sequence data at this locus is available for *S. oleoides* at GenBank. *trnL-F* spacer region is also routinely employed for phylogenetic and phylogeographic studies of angiosperms.

## Materials and Methods

### 1. Taxon Sampling

Young leaves were collected from 20 geographic isolates of *Salvadora oleoides* covering the greater part of its range in the Indian subcontinent (Table 1 and Fig 1). Leaf samples from three individual trees from each location were separately analyzed, including PCR and sequencing at two loci, for intra-population heterogeneity. No special permission was required to collect these plant materials, as none of the sampling locations were part of the areas designated as protected by Government of Punjab. Collected specimens were transported to the laboratory in zip-lock polythene bags. Pressed vouchers were prepared and deposited in the Central National Herbarium, Botanical Survey of India, Calcutta (*Index Herbariorum* Code: CAL). Samples for molecular analysis were stored at – 80°C till further analysis.

**Table 1.** Details of geographical isolates analyzed in this study. A dash (-) indicate no data available.

| Voucher No. | Location | Collected by | Collection date | Latitude (N) | Longitude (E) | ITS1-5.8S-ITS2 sequence generated (GenBank Accession) | *trnL-F* spacer sequence generated (GenBank Accession) |
| --- | --- | --- | --- | --- | --- | --- | --- |
| CUP VOUCHER-SO-2014-1 | Faridkot | Sukhwant Singh and Navreet Kaur | 27-02-2014 | 30°38'05.62''N | 74°47'28.10''E | KP795947 | KP795970 |
| CUP VOUCHER-SO-2014-2 | Sangrur | Sukhwant Singh and Navreet Kaur | 02-03-2014 | 30°15'22.61''N | 75°44'42.88''E | KP795949 | KP795958 |
| CUP VOUCHER-SO-2014-3 | Mansa | Sukhwant Singh and Navreet Kaur | 07-03-2014 | 30°06'18.20''N | 75°10'39.55''E | - | KP795969 |
| CUP VOUCHER-SO-2014-4 | Talwandi Sabo | Sukhwant Singh and Navreet Kaur | 07-03-2014 | 29°59'48.65''N | 75°09'32.08''E | KP795946 | KP795965 |
| CUP VOUCHER-SO-2014-5 | Barnala | Sukhwant Singh and Navreet Kaur | 08-03-2014 | 30°19'35.18''N | 75°30'10.70''E | KP795950 | KP795968 |
| CUP<br>VOUCHER-SO-2014-6 | Patiala | Sukhwant Singh<br>and Navreet<br>Kaur | 08-03-2014 | 30°16'24.70''N | 75°58'35.37''E | - | KP795964 |
| CUP<br>VOUCHER-SO-2014-7 | Taran Taran | Gurbeer Singh | 15-03-2014 | 31°10'46.35''N | 74°54'12.08''E | - | KP795962 |
| CUP<br>VOUCHER-SO-2014-8 | Malaut | Jasdeep Singh | 18-03-2014 | 30°11'36.31''N | 74°24'33.11''E | - | KP795961 |
| CUP<br>VOUCHER-SO-2014-9 | Moga | Sukhwant Singh | 02-04-2014 | 30°43'10.13''N | 75°06'02.04''E | - | KP795960 |
| CUP<br>VOUCHER-SO-2014-10 | Hoshiarpur | Gurpreet Singh | 03-04-2014 | 31°38'08.66''N | 75°50'14.66''E | - | - |
| CUP<br>VOUCHER-SO-2014-11 | Ferozpur | Sukhwant Singh<br>and Navreet<br>Kaur | 04-04-2014 | 30°53'38.79''N | 74°56'23.46''E | - | KP795967 |
| CUP<br>VOUCHER-SO-2014-12 | Amritsar | Sukhwant Singh | 07-04-2014 | 31°38'09.89''N | 74°49'27.79''E | - | - |
| CUP<br>VOUCHER-SO-2014-13 | Zira | Sukhwant Singh<br>and Navreet<br>Kaur | 09-04-2014 | 30°52'43.38''N | 74°59'16.70''E | KP795951 | KP795963 |
| CUP<br>VOUCHER-SO-2014-14 | Kotkapura | Sukhwant Singh<br>and Navreet<br>Kaur | 09-05-2014 | 30°38'20.34''N | 74°49'46.35''E | KP795955 | - |
| CUP<br>VOUCHER-SO-2014-15 | Bathinda | Navreet Kaur | 21-05-2014 | 30°11'45.09''N | 74°56'42.99''E | KP795948 | KP795957 |
| CUP<br>VOUCHER-SO-2014-16 | Chandigarh | Sukhwant Singh<br>and Navreet<br>Kaur | 22-05-2014 | 30°46'37.19''N | 76°45'34.70''E | - | - |
| CUP<br>VOUCHER-SO-2014-17 | Jalandhar | Sukhwant Singh<br>and Sukhdeep<br>Kaur | 03-06-2014 | 31°07'91.78''N | 75°33'57.29''E | KP795952 | - |
| CUP<br>VOUCHER-SO-2014-18 | Ludhiana | Sukhwant Singh | 06-06-2014 | 30°67'49.13''N | 75°45'74.08''E | KP795956 | - |
| CUP<br>VOUCHER-SO-2014-19 | Abohar | Harjoban Singh | 12-06-2014 | 30°04'09.80''N | 74°19'21.23''E | KP795953 | KP795966 |
| CUP<br>VOUCHER-SO-2014-20 | Muktsar | Harpreet Singh | 13-06-2014 | 30°40'89.56''N | 74°52'20.13''E | KP795954 | KP795959 |

**Figure 1.**
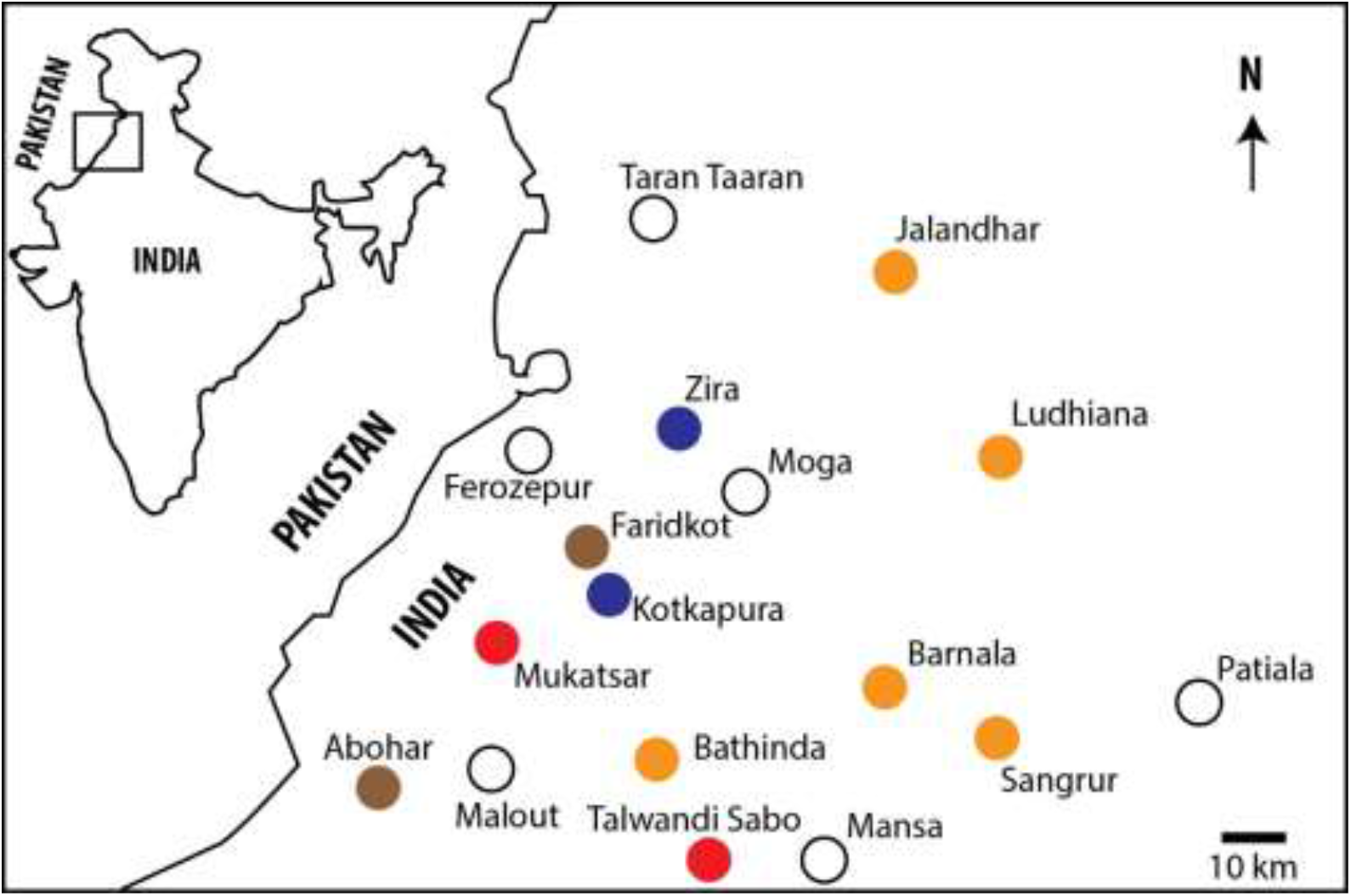
Map indicating sampling locations. ITS haplotypes shown in Fig 2 are represented in colored icons. Red = Clade 1; Blue = Clade 2; Brown = clade 3; and Dark Brown = clade 4. Empty circles represent other locations included in the present study. Zoomed location in Indian map is highlighted in Inset.

### 2. DNA Extraction and PCR Amplification

Total genomic DNA was extracted from the specimens using Hi PurA^TM^ Plant Genomic Extraction Kit (Hi Media Laboratories Pvt. Ltd., Mumbai) following manufacturer’s protocol. The quality of DNA was checked on 0.8% agarose gel, and the quantity of DNA was determined with a spectrophotometer (NanoDrop2000, Thermo Scientific). Extracted DNA was stored at – 20°C. For PCR amplification, 20μl reaction mixture was prepared containing 2μl of 10X reaction buffer with 1.8μl 3mM magnesium chloride (Applied Biosystems, Foster City, CA, USA), 4μl each of 10 μm primers: ITS1 (5’ TCCGTAGGTGAACCTGCGG 3’)(White et al. 1990), ITS4 (5’ TCCTCCGCTTATTGATATGC 3’) (White et al. 1990), *trnL-F* (5’ CGAAATCGGTAGACGCTACG 3’) (Poeaim et al. 2012), *trnL-R* (5’ ATTTGAACTGGTGACACGAG 3’) (Baraket et al. 2008), 2μl of 10μm dNTP mixture containing dATP, dTTP, dCTP and dGTP (Genaxy Scientific, India), 0.2μl of 5u/500μl Taq DNA Polymerase (Thermo Scientific, India), 4μl of template DNA and 2μl sterile water. PCR amplification were carried out in a programmable thermal cycler (Bio-Rad, India) and reaction profile included an initial denaturation at 94°C for five minutes, followed by 35 cycles of 94°C for 1 minute, 52°C for 2 minutes and 72°C for 2 minutes and a final extension of 72°C for 10 minutes. Amplified products were electrophoresed on 1.5% agarose gel for 25 minutes at 250V and visualized with ethidium bromide in order to determine approximate length and purity.

### 3. Purification of PCR Product and DNA Sequencing

Positive reactions were purified using Exo SAP-IT PCR clean-up kit following manufacturer’s instructions (USB Corporation, Cleveland, OH, USA). The purified PCR products were subjected to chain termination reaction with the following composition of reaction mixture – buffer 2μl, sequence-ready reaction mixture 1μl, primer 2μl, PCR product 2μl and autoclaved distilled water 3μl. The PCR conditions for chain termination reaction were as follows – 95°C for 5 minutes, followed by 50 cycles of denaturation at 95°C for 45 seconds, annealing at 52°C for 30 seconds and extension at 60°C for 4 minutes. For cleaning of sequencing reaction, Big-Dye X terminator kit (Applied Biosystems, Foster City, CA, USA) was used, following manufacturer’s protocol. Purified PCR products were subjected to bidirectional Sanger sequencing using a dideoxy chain termination protocol with ABI BigDye Terminator Cycle Sequencing Ready Reaction Kit v3.1 (Applied Biosystems, Foster City, CA, USA) and a programmable thermal cycler (Veriti, ABI, USA), as per (Bast et al. 2014). DNA sequences were assembled using the computer program CodonCodeAligner (CodonCode Corporation, USA). Out of 20 isolates, ITS1-5.8S-ITS2 locus from only 11 isolates got amplified and its sequence length ranged between 605-658 bp. *TrnL-F* spacer locus from 14 isolates got amplified and its sequence length ranged between 188-478 bp. Out of 75 generated sequences, only 25 were unique, as three individual trees at each of the sampling populations (geographical locations) had exactly same sequences for both of the genomic loci analyzed. Unique 25 sequences were deposited in Genbank (Table 1).

## Multiple Alignment and Phylogenetic Analyses

Step-by-step protocol followed for contig assembly, multiple sequence alignment, ML ModelTest, and phylogenetic analyzes are available at Nature Protocols (Bast 2013). In summary, sequences were assembled using CodonCode Aligner, BLASTN sequence homology search, Multiple Sequence Alignments, consensus sequence generation, and Bayesian Inference (BI) phylogenetic inference (using MrBayes add-in) were conducted in Geneious Pro (Biomatters, NZ), and Model Selection, pairwise distance estimation, and Maximum Likelihood phylogenetic inference were conducted in MEGA. For intraspecific phylogeographic assessments, an accession of *Khaya nyasica* (Sapindales) was used as an outgroup for ITS dataset and an accession of *Borthwickia trifoliata* (Brassicales) for *trnL-F* spacer dataset. For phylogenetic assessment of malvids, we used accessions of *Euphorbia* spp. as outgroups, as in (Rodman et al. 1998). The rationale for choosing these outgroups was to have taxa sufficiently closely related to the ingroup for resolving phylogenetic structures to the maximum extent. For example, as Malvid gene tree based on ITS revealed a phylogenetic affinity of *Salvadora* to Sapindales, we used another member of Sapindales rather than Brassicales as an outgroup for intraspecific phylogeographic assessment of *Salvadora oleoides*. We have concurrently tested a number of different outgroup taxa, including as far as *Ficus* (Moraceae), but the topology of phylogram remained similar indicating the robustness of the method.

## Results

### Intra-population Heterogeneity

Neither within-individual polymorphisms (dual-peaks at electropherograms/heterozygosity due to imperfect homogenization of multiple rRNA genes, paralogs, pseudo-genes, etc.) nor within-population polymorphisms were observed at either of the two loci analyzed. Three sequences at each of the sampling populations had no nucleotide differences either at ITS1-5.8S-ITS2, or at *trnL-F* spacer region.

## BLASTN

BLASTN Homology search with NCBI nr nucleotide database (word size = 10, Max E-value =1e-1 and gap cost-open extend = 5.2) revealed interesting results. ITS sequences, both individually, as well as the consensus sequences generated for 11 of our ingroup, isolates, retrieved *Arfeuillea arborescens* (EU720461) as the closest hit with an E-value of 3.24e-93 and 79% identity. Particulars of top 10 hits are summarized in Table 2, and all 10 of them were from order Sapindales. This came as a surprise, as the current consensus among molecular phylogeneticists working on angiosperm systematics is that the family Salvadoraceae belongs to Brassicales. Top 100 results included orders as diverse as Fagales (*Nothofagus*) and Malpighiales (*Quiina, Lacunaria*) however not a single accession belonging to Brassicales. It was not because of the non-existence of ITS sequences of Brassicales in GenBank; as of this writing there are 5990 sequences of ITS locus belong to the members of Brassicales exist in the database. *TrnL-F* spacer sequences, both individually, as well as the consensus sequences generated for 14 of our ingroup isolates, retrieved *Salvadora angustifolia* (KC479309) as the closest hit with an E-value of 0 and 99.4% identity, well within generally accepted conspecific threshold (98.7%, (Schloss and Handelsman 2004)). Particulars of top 10 hits were summarized in Table 3, and all 10 of them were from order Brassicales. Comparable results were obtained in discontinuous Mega BLAST as well (results not presented), with all top ten hits of ITS belonging to Sapindales, and that of *trnL-F* spacer belong to Brassicales.

**Table 2.** Particulars of top 10 BLASTN hits of consensus sequence of ITS region from *Salvadora oleoides* isolates from Punjab, India. All ten were from order Sapindales.

| S. No | Species | GenBank Accession | Percent identity |
| --- | --- | --- | --- |
| 1 | *Arfeuillea arborescens* | EU720461 | 79 |
| 2 | *Acer laurinum* | AM113543 | 80 |
| 3 | *Acer laurinum* | AM113542 | 79 |
| 4 | *Protium serratum* | KJ503508 | 80 |
| 5 | *Protium cranipyrenum* | KJ503531 | 80 |
| 6 | *Protium ravenii* | KJ503448 | 80 |
| 7 | *Atalaya alata* | EU720425 | 79 |
| 8 | *Protium glabrescens* | KJ503444 | 79 |
| 9 | *Protium urophyllidium* | AY375516 | 80 |
| 10 | *Acer laurinum* | DQ366114 | 79 |

**Table 3.**
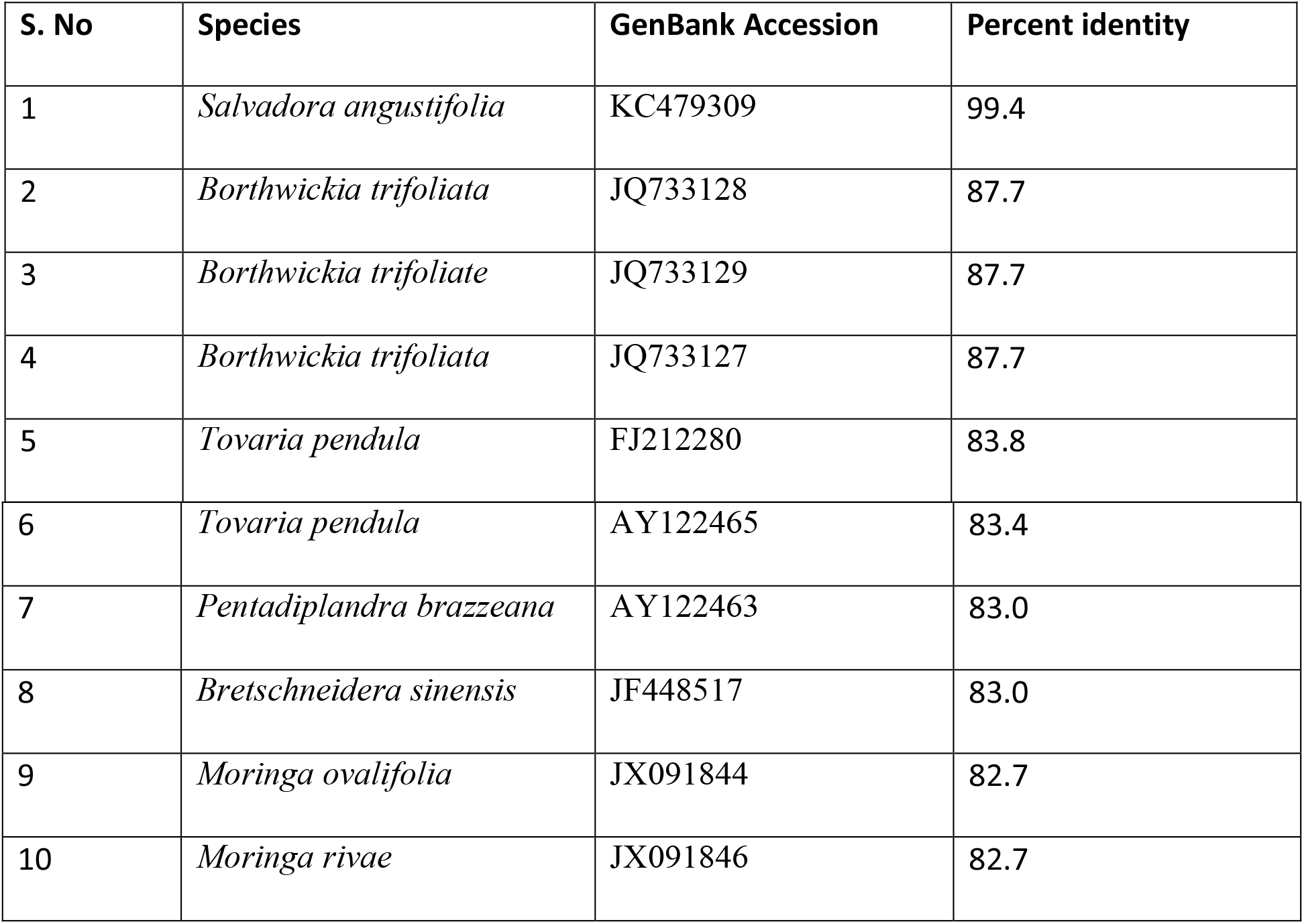
Particulars of top 10 BLASTN hits of consensus sequence of *trnL-F* spacer region from *Salvadora oleoides* isolates from Punjab, India. All ten were from order Brassicales.

## Phylogeography

Results of phylogeographic assessment using ITS and *trnL-F* spacer regions are presented as Fig. 2. In summary, ITS phylogram resolved intraspecific clades better than *trnL-F* spacer phylogram. Intraspecific genetic diversity (within-group mean distance) for ITS dataset (0.020) was found to be higher than that of *trnL-F* spacer dataset (0.003). Corrected (T92 model) pairwise distances of ITS dataset ranged between 0, between isolates Jalandhar and Ludhiana, and 0.063 between isolates Zira and Talwandi Sabo. Corrected (JC69 model) pairwise distances of *trnL-F* dataset ranged between 0, between isolates Moga and Zira, and 0.008 between isolates Barnala and Zira.

**Figure 2.**
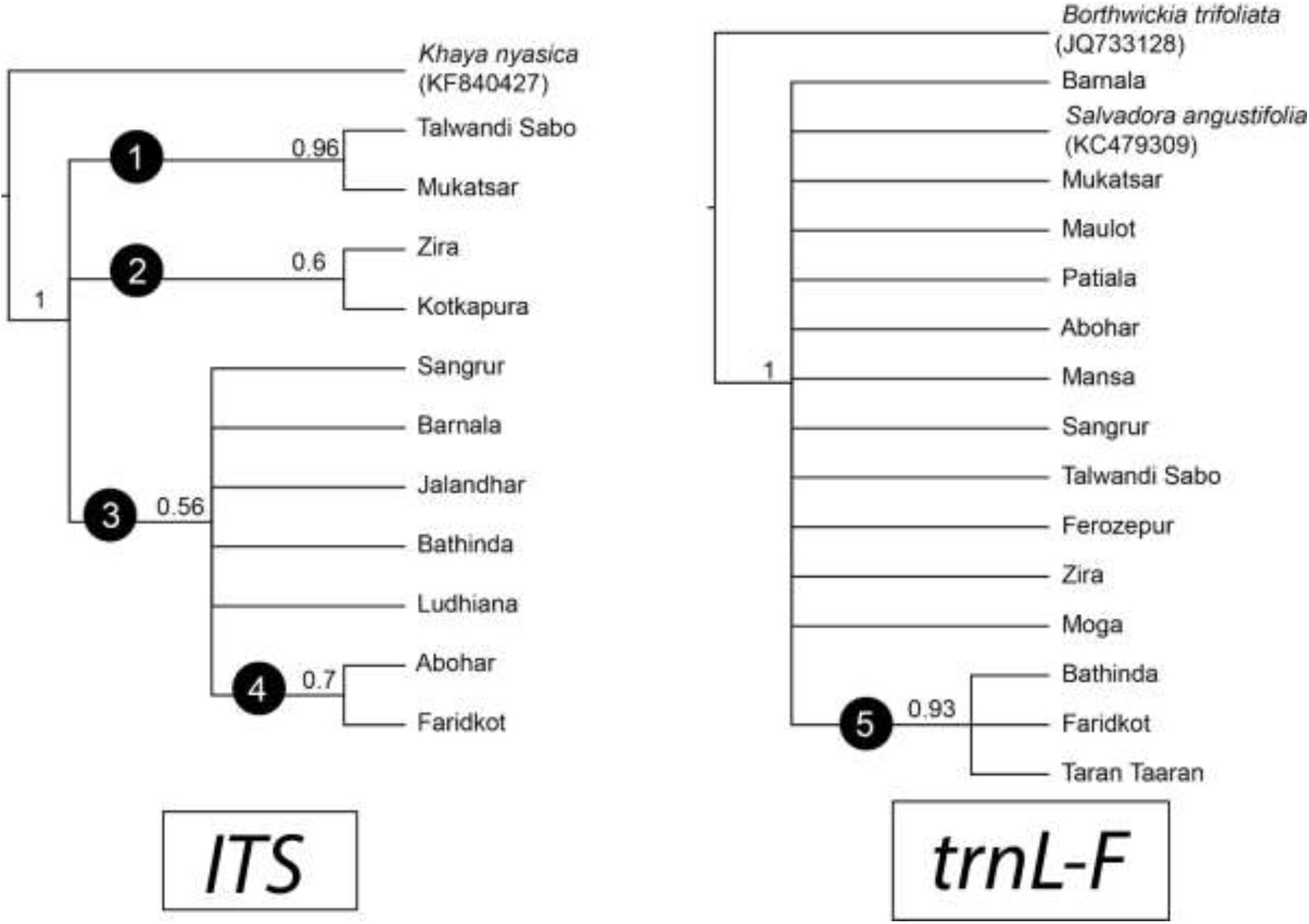
Bayesian Inference (BI) Molecular Phylogeny inferred from ITS (left) and *trnL-F* (right) sequences. The evolutionary history was inferred by using the Bayesian Inference method based on the JC69 (for ITS) and T92 (for *trnL-F*) models. For ITS tree, mean LnL = – 1642.375 and effective sample size = 4059.229. For *trnL-F* tree, mean LnL = – 1005.345 and effective sample size = 3504.021.

ITS phylogram resolved four clades. In general, geographically closer isolates clustered into distinct clades. The best-supported clade (Clade 1) consisted of isolates from Mukatsar and Talwandi Sabo, followed by a clade consisting of Abohar and Faridkot (Clade 4). The later clade (clade 4) nested within a larger clade consisting of the majority of sampled isolates (clade 3). Another, moderately supported, clade consisted of isolates from Kotkapura and Zira (clade 2). Clades 1, 2 and 3 formed a trichotomy, with each clade being sister to the rest two clades. *trnL-F* phylogram resolved only one clade (clade 5) that consisted of isolates from Bathinda, Faridkot, and Taran Taaran. All other isolates occupied a basal position in a giant polytomy, perhaps indicating low phylogenetic signal for sufficient resolution. Interestingly, an accession of *Salvadora angustifolia* (KC479309) clustered within this polytomy. The *trnL-F* phylogram broach few possibilities; either a cryptic speciation inside the range of parent population (as *Salvadora angustifolia* clustered within the clade), or synonymy of *Salvadora oleoides* and *Salvadora angustifolia* (as the group consisting of these two taxa was polyphyletic). Considering clades in ITS phylogram as haplotype variants, respective geographical locations of these haplotypes are labeled in Fig. 1.

## Phylogenetic position of Salvadorales within Malvids

Results of our ITS sequence-based phylogenetic assessment of the comparative position of family Salvadoraceae within malvids corroborated our earlier observation of the BLASTN affinity of this family with Sapindales (Fig 3). *Salvadora* clustered within the strongly supported monophyletic clade of Sapindales and occupied a position phylogenetically closer to family meliaceae. However, our *trnL-F* spacer-based phylogram clustered Salvadoraceae within Brassicales with strong bootstrap support (Fig. 4).

**Figure 3.**
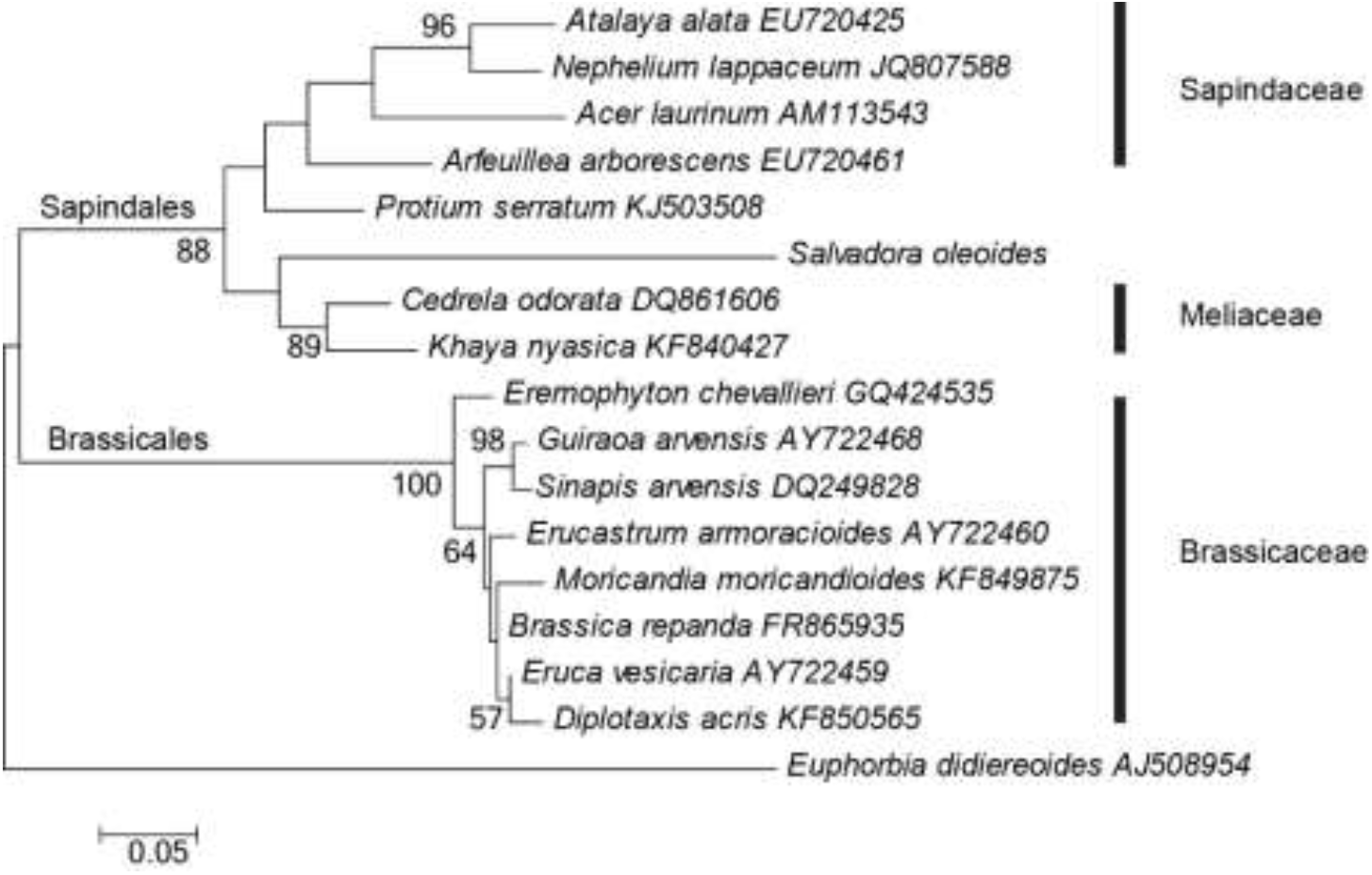
Maximum likelihood (ML) phylogram based on ITS1-5.8S-ITS2 sequences using K2P+G model of molecular evolution in MEGA phylogenetic framework. Numbers near nodes represent ML bootstrap proportion exceeding 50. LnL = −1860.2630. Bootstrap proportions (500 replicates) exceeding 50 were shown near nodes. Scale bar is on the unit of average nucleotide substitutions per site. The analysis involved 17 nucleotide sequences. All positions with less than 95% site coverage were eliminated. That is, fewer than 5% alignment gaps, missing data, and ambiguous bases were allowed at any position. There were a total of 335 positions in the final dataset.

**Figure 4.**
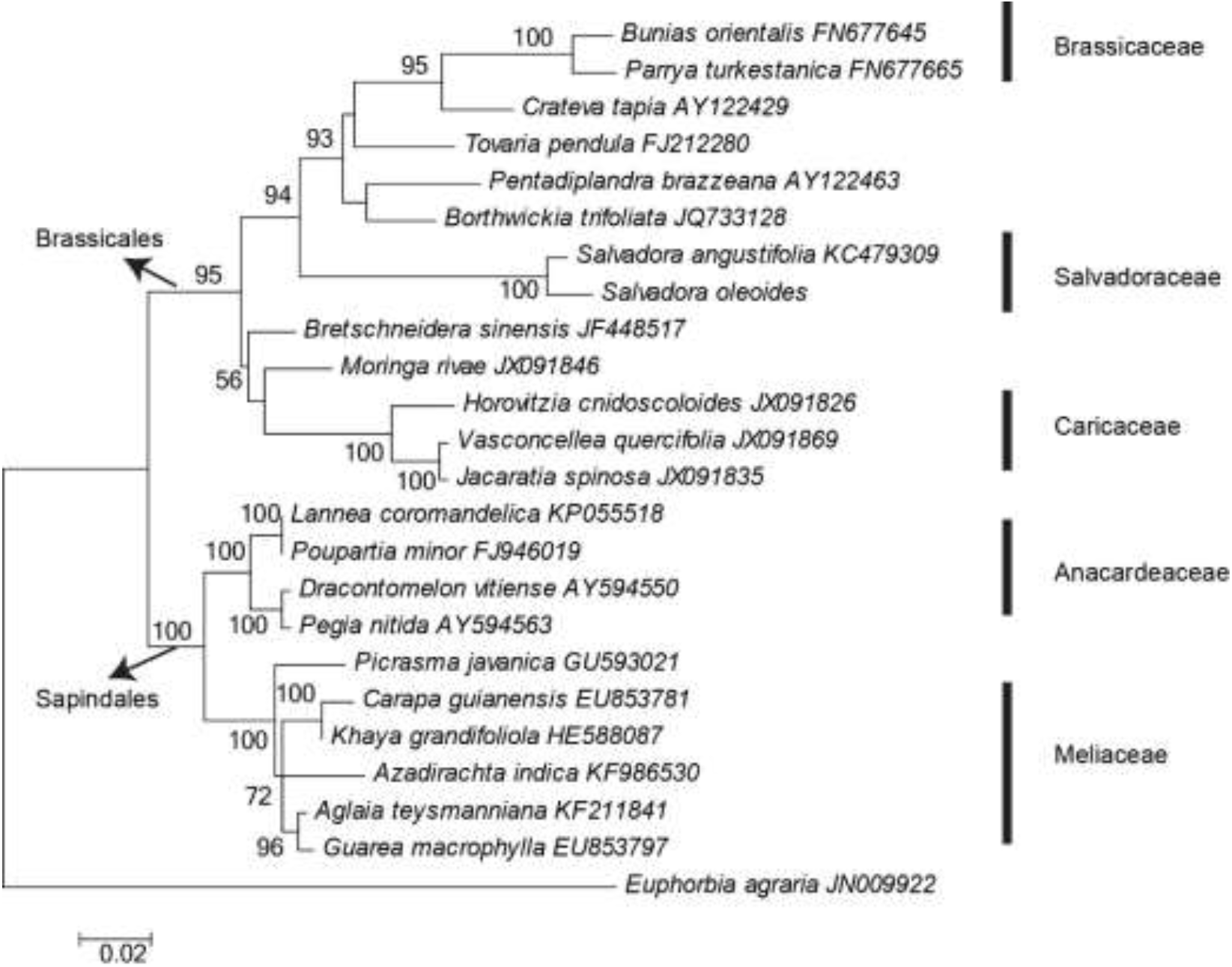
Maximum likelihood (ML) phylogram based on *trnL-F* sequences using T92+G model of molecular evolution in MEGA phylogenetic framework. Numbers near nodes represent ML bootstrap proportion exceeding 50. LnL = −2302.8388. Bootstrap proportions (500 replicates) exceeding 50 were shown near nodes. Scale bar is on the unit of average nucleotide substitutions per site. The analysis involved 24 nucleotide sequences. All positions with less than 95% site coverage were eliminated. That is, fewer than 5% alignment gaps, missing data, and ambiguous bases were allowed at any position. There were a total of 445 positions in the final dataset.

## Discussion

This study generated DNA barcode data for *Salvadora oleoides* for the first time in the world. Comparing with *trnL-F*, ITS locus exhibited more within-group distance, as well as resolved finer phylogeographic structures. It can be concluded that ITS is a useful locus for within species phylogeographic assessment in this species. Most probably this is due to differential evolutionary rates for these two loci; ITS-being an intron within the nucleoribosomal region‐ is more prone to mutations, while *trnL-F* being an intergenic spacer is more evolutionarily conserved. On the other hand, in phylogenetic analysis within malvids using ITS, a longer branch-length observed for the consensus sequence of our isolates suggest that it is a rapidly evolving Operational Taxonomic Unit with elevated substitution rates. Indeed, long stretch of insertions (22 nucleotides long) are observed for our consensus sequence near the end of ITS2 region, between sites 636 and 658, in the ITS alignment. As the method we adopted to infer the phylogeny was Maximum Likelihood-which had been proven to be less prone to Long Branch Artifacts, it is likely that this disproportionately long branch length was not due to any artifacts. One likely explanation is allopatric speciation. Within-group mean distance observed for our isolates in ITS dataset (0.02) is indeed the highest ever recorded among angiosperms. Given that the isolates are sampled from area spanning ca 23000 km^2^ with no apparent physical barriers between them, barriers in the form of anthropogenic islands due to habitat destruction could very well construe a vicariance mediator and likelihood for such a scenario of allopatric speciation is quite high. A word of caution in this conclusion is deemed necessary. While the current patterns of geographic differentiation may ultimately lead to long-term divergence and speciation, these might also be latest in several cycles of ephemeral patterns of genetic differentiation that this long-lived species have experienced.

Our phylogeographic assessment revealed the existence of five intraspecific clades of *S. oleoides*. Although the correlation between evolutionary distances and genetic distances were rather weakly negative (i.e., geographically close isolated had larger genetic distances between them, *R*^2^ = 0.0211), geographically closely located isolates were part of distinct clades in our phylograms. This observation in turn suggests an evolutionary congruence across geographic space. As DNA barcode data for this species is non-existent in public sequence repositories prior to this study, we were unable to perform finer phylogeographic assessments, including its center of origin and dispersal routes. It can be nevertheless concluded that factors such as habitat destruction through agricultural expansion of plains might have forced the remaining population of this tree to isolated patches of thorn forest biotas analogous to an archipelago. Our analyses were insufficient to conclude whether vicariance phenomena or range expansion through geodispersal shaped the current patterns of its distribution. Separation of once contiguous area of species distribution, as observed in this study, do not warrant these endemic spots are related geologically. We can only infer that these biotas were once part of a continuous range and most probable explanation for the apparent vicariance is the anthropogenic agricultural expansion.

An important aspect of our *trnL-F* phylogram is that it clustered *S. angustifolia* within our geographical isolates of *S. oleoides*. In addition, the percent identity between the consensus sequence of *S. oleoides* and *S. angustifolia*, 99.4%, was well within the intraspecific range commonly accepted for angiosperms. It can be inferred that these two species are conspecific. According to the principle of priority (McNeill 2012) the binomen *S. oleoides* (Decne, 1844) should be preferred over *S. angustifolia* (Turril, 1918).

The most surprising finding is the affinity of salvadoraceae in Sapindales as revealed by our phylogram based on ITS locus. In our knowledge, this is the first study that revealed such an affinity using molecular phylogeny. This finding corroborates an earlier morphology-based systematic scheme that grouped Salvadoraceae under Sapindales. However, as per our *TrnL-F* phylogram, Salvadoraceae is affiliated in Brassicales, as currently considered by the majority of angiosperm systematicists. The reason for this contrasting observation remain elusive; this might be related to the differential manner in which these two loci evolve along plant lineages. ITS, being a nuclear intron, are prone to recombination and follows biparental inheritance. *trnL-F*, on the other hand, being an intergenic spacer in plastid genome, are not prone to recombination and follows the typical matrilineal pattern of inheritance. However, phylogeny based on 18S locus, another nuclear region contiguous to the ITS intron, revealed affiliation of Salvadoraceae in Brassicales (Rodman et al. 1998). Our conflicting finding warrants need for further scrutiny of angiosperm phylogeny based on multi-local concatenated approach with integrated morphometric dataset, for having the total evidence [*supermatrix*, (Sanderson 1998)] at our disposal. In earlier molecular systematic analyzes as well as comparative analyzes based on floral anatomy (De Craene and Wanntorp 2009) and seed anatomy (Tobe and Raven 2012), Salvadoraceae had been shown to be evolutionarily closely related with Bataceae and Koberliniaceae. Although we were really interested in this affinity, as well as relative phylogenetic positions of families Bataceae and Koberliniaceae within malvids, neither ITS nor *trnL-F* sequences from any of its members are currently available at GenBank. A few other observations in our phylogenetic analysis of malvids are deemed noteworthy. Our ITS phylogram revealed non-monophyly of family sapindaceae, with *Acer*, a member of aceraceae, clustered within the clade consisting otherwise of the members of sapindaceae. As per our *trnL-F* phylogram, affiliation of *Picrasma*, currently considered to be a member of family Picrasmaceae, within the family Meliaceae was obvious, as meliaceae + *Picrasma* had a strongly supported (BP = 100) clade. Similarly, the genus *Crateva*, currently considered to be part of family cratevaceae, showed strong affinity (BP = 95) to the family Brassicaceae.

This is the first DNA-sequence based phylogeographic assessment of *Salvadora* in the world. Sequence based phylogeographic assessment of tropical tree species is a relative rarity and in India, such investigations are almost non-existent. It is hopeful that the natural history of *S. oleoides* in Indian Subcontinent revealed in the present study would be beneficial for further, larger-scale phylogeographic assessment of this species for testing various hypotheses on geological, climatological and anthropogenic factors that might have shaped its current distribution. In addition, it is hopeful that our revelation of the affinity of Salvadoraceae to Sapindales would foster further, more detailed, “total evidence” assessment of malvids by molecular systematicists working on this field.

## Acknowledgements

We thank Dr. Kenneth J. Sytsma, University of Wisconsin, Madison for helpful insights. We are thankful to Dr. Pankaj Bhardwaj, Ms. Pooja Rani and Ms. Sheetal Sharma for support with experimental works. We are also thankful for the Vice-chancellor, the Central University of Punjab for his support with respect to the execution of this research. The study was supported by grant-in-aid “Research Seed Money” scheme of the Central University of Punjab.

## Author’s Contributions

FB conceived the topic, interpreted the data and drafted the manuscript. NK collected the samples and performed the experiments.

## Funding

The study was funded by a grant from Research Seed Money programme of Central University of Punjab, India

## Conflict of Interest

The authors declare that they have no conflict of interest.

